# Designing membrane-reshaping nanostructures through artificial evolution

**DOI:** 10.1101/2020.02.27.968149

**Authors:** Joel C. Forster, Johannes Krausser, Manish R. Vuyyuru, Buzz Baum, Anđela Šarić

## Abstract

In this paper we combine the rules of natural evolution with molecular dynamics simulations to design a nanostructure with a desired function. We apply this scheme to the case of a ligand-covered nanoparticle and evolve ligand patterns that promote efficient cell uptake. Surprisingly, we find that in the regime of low ligand number the fittest structures are characterised by ligands arranged into long one-dimensional chains that pattern the surface of the particle. We show that these chains of ligands provide particles with high rotational freedom and they lower the free energy barrier for membrane crossing. This demonstrates the efficacy of artificial evolution to identify non-intuitive design rules and reveals a new principle of design that can be used to inform artificial nanoparticle construction and the search for inhibitors of viral entry.

---

The mechanistic rules that govern the function of the nanoscale machinery of life remain poorly understood. Our intuition for how nanostructures in nature operate is guided by experimental observation interpreted through mechanistic models. Such a strategy rests on the researchers’ ability to navigate the large phase space of possible model ingredients and parameters to capture the key physics behind complex biological phenomena.

Here, we set out to take a reverse approach: instead of deducing the design rules of nanostructures by observing nature, we specify the function that the nanostructure should perform and use a computer model to evolve its design. To this effect we couple the principles of biological evolution to molecular dynamics (MD) simulations. The computational protocol starts with a random population of nanostructure designs and measures how well the individual structures perform in MD simulations, as shown in the schematic in Fig. 1 (a). We then use an evolutionary algorithm (GA) to mutate and breed the fittest members of the population and repeat the process to evolve better designs. The MD-GA procedure is iterated until the nanostructures generated by rounds of mutation and selection can perform the task efficiently. By observing the structures as their performance evolves, we can identify the key features needed for a desired function. This reverse approach can be used to identify engineering principles that we would not have a priori postulated based purely on intuition. Thus it represents an orthogonal way of exploring the physical mechanisms of nanostructure design.

**Fig. 1.**
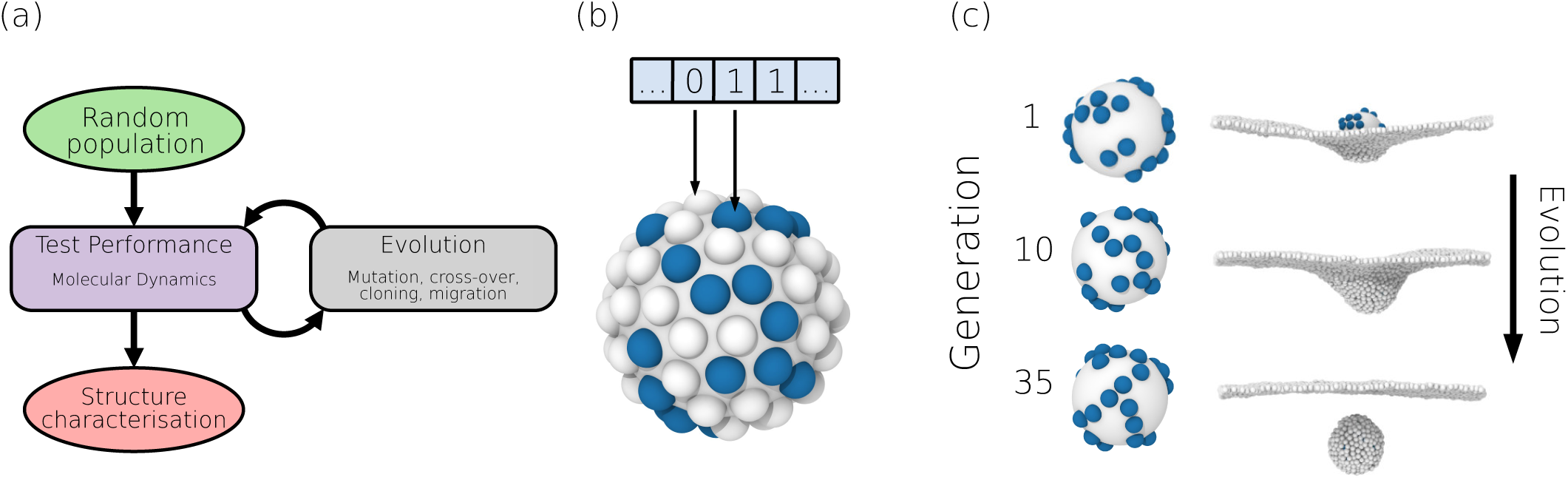
Combining molecular dynamics and a genetic algorithm. (a) Graphical representation of the MD-GA scheme developed here. Simulations are used to asses the performance of the nanostructure designs, while the evolutionary algorithm is used to optimise the designs towards an ever better performance. At the end of the proccess the evolved designs were structurally characterised. (b) Implementation of the nanoparticle design: the nanoparticle is covered by 72 ligand, of which *N* are active and can bind the membrane. The active ligands are coloured in blue and are represented as “1” in the nanoparticle genome. (c) Exemplary designs from three different generations along with a snapshot from the last timestep of their respective simulation runs. After performing a series of MD-GA iterations that reposition the active ligands on the nanoparticle surface, while keeping their number and interaction strength constant throughout evolution, the nanoparticle designs improve and the particles become more successful in crossing the membrane.

As a proof of principle, we apply the framework to study the ability of a ligand-covered nanoparticle to cross a membrane. In this way we determine the optimal ligand pattern for this task. We show that this approach identified non-trivial nanoparticle designs, whose kinetic and thermodynamics features we have analysed to determine how they drive efficient particle internalisation.

## Simulation Model

The nanoparticle is modelled as a rigid body, made up of a central particle that carries ligands on its surface to facilitate binding to a fluid membrane, as shown in Fig. 1 (b). To curve the membrane and form a vesicle (1, 2) a minimal interaction strength between the ligands and the membrane is required. Importantly, for a given interaction strength between ligands and the membrane, the efficiency of membrane crossing depends on the arrangement of ligands on the nanoparticle. For instance, in the limiting case where all the ligands are clustered at one pole, the nanoparticle binds strongly to the membrane, but cannot be wrapped by the membrane and internalised.

To account for different ligand arrangements the nanoparticle was fully covered by 72 ligand sites, while *N* of those are active sites, which are able to bind to the membrane beads via a generic Lennard-Jones potential of a depth *E*, see Fig. 1 (b) and Methods. The membrane was modelled using a single particle-thick model (3), which reproduces the correct mechanical properties of biological membranes and is capable of fusion and fission. Simulations were run using Langevin dynamics within the LAMMPS molecular dynamics package (4). For more details see Methods.

## Evolutionary Algorithm

To explore efficient nanoparticle designs, we kept the total number of active ligands and their binding strength to the membrane constant throughout evolution. The ligand design was turned into a 1D single bit arrays, as illustrated in Fig. 1 (b), which enabled the following genetic algorithm operations: tournament selection, two point crossover, and shuffling mutation. To avoid premature fixation, the population was split into independent “demes” and permitted to trade individuals every generation (5, 6).

As a measurement of the “fitness” of the membrane-crossing nanoparticle we used the degree of wrapping by the membrane. Budding is typically achieved after a particle binds, become wrapped by membrane, and detaches from the mother membrane. The fitness function used to analyse the performance of a particle is defined as:

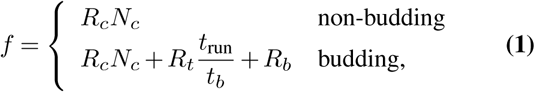

where *N*_*c*_ is the number of membrane beads in contact with the ligands and is assigned a weighting constant *R*_*c*_. When budding occurred an additional reward, *R*_*b*_, was given. To characterise the efficacy of budding we measured the time of budding *t*_*b*_, normalised by the runtime of the total simulation *t*_run_, which is equal for all the simulations, and assigned it a weighting constant *R*_*t*_. Further details are given in Methods. The genetic algorithm was run for 35 generations across 4 demes, each with a population of 20 individual particles. Each individual particle *i* was run for 4 different initial rotational positions *k* to ensure the robustness of particle uptake. The corresponding fitness *f*_*k*_ was calculated for each run and averaged to obtain the mean particle fitness of the particle 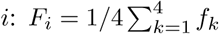. The average population fitness for a single generation is then 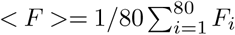.

This was repeated for different number of active ligands (19 *< N <* 61) and ligand-membrane attraction strengths (3*kT < ϵ <* 15*kT*), producing a total population of around 1.298 × 10^6^ individuals, with 5.34 × 10^5^ being unique. A full overview of all major steps involved in the algorithm is provided in the Supplementary Information, Fig. S1.

## Nanoparticle evolution

An example of a typical nanoparticle evolution is shown in Fig. 1 (c). The random nanoparticle design in the first generation is not able to penetrate the membrane at the end of the MD run, whereas the evolved designs deform the membrane more strongly, and eventually are able to form a vesicle and bud off freely on the other side of the membrane.

Accordingly, the mean fitness of the whole population *< F >* increases as the evolution progresses and eventually approaches saturation, as shown in Fig. 2 (a) for three different combinations of the ligand number *N* and ligand-membrane interaction strengths *ε*. The fitness values are normalised by the mean fitness value of the population of randomly generated particles, showing that the evolved particles outperform the randomly generated ones. The maximum fitness that the population is however able to reach depends on the combination of the number and arrangement of ligands and their affinity.

**Fig. 2.**
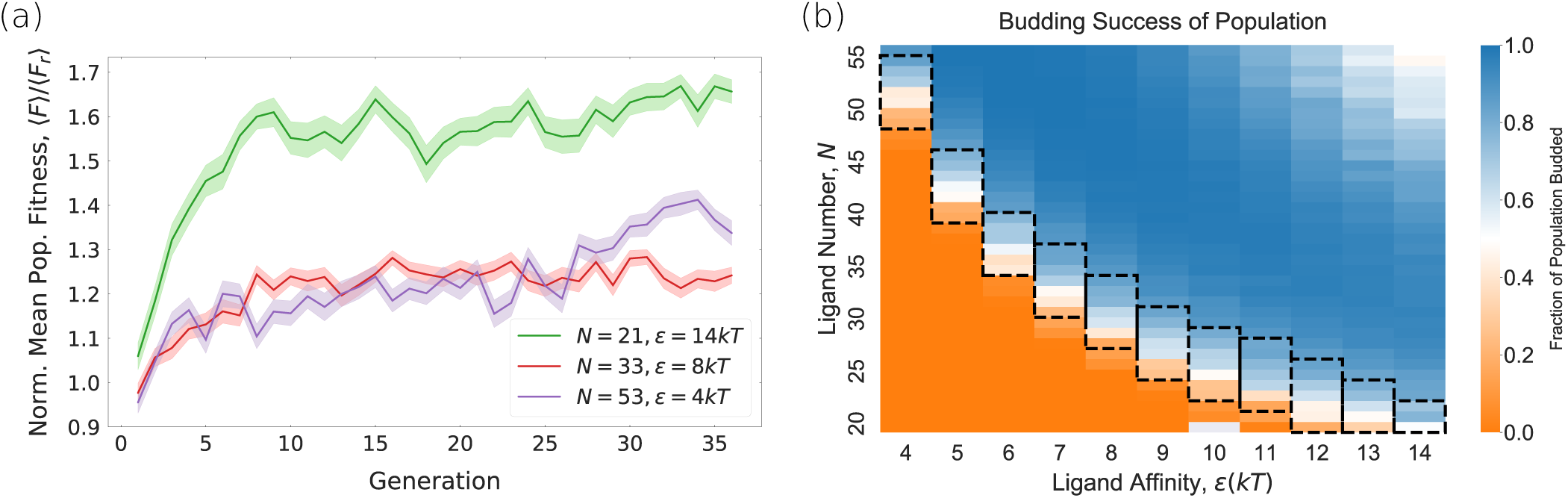
Performance of the evolved nanoparticles. (a) The mean population fitness *< F >*, normalised by the mean fitness of a random population of particles with the same *ϵ* and *N*, *< F*_*r*_ *>*. These curves measure how much the particle is wrapped by the membrane and how quickly it crosses it (Eq. (1)), shown here for three examples of the active ligand number *N* and ligand-membrane interaction strength *ϵ*. The fitness value increases as the evolution proceeds for all populations. (b) The fraction of the entire population that successfully crossed the membrane at various values of the active of ligands *N* and ligand-membrane binding energy *ϵ*. In the region around *N*_*ϵ*_ *≈* 200*kT* (marked by black boxes) the population is made up of a mixed collection of budding and non-budding particles; here the nanoparticle design plays a crucial role.

To better understand the effect of the ligand distribution on the nanoparticle’s ability to traverse through the membrane we measure the average budding success at different number of ligands and ligand-membrane interaction strengths across the whole GA evolution. For large numbers of active ligands and large binding strengths almost every design will traverse the membrane successfully in the first generation, as shown in Fig. 2 (b). In this regime the ligand arrangement is not the deciding factor, since large *N* entail a total adhesion energy large enough to induce membrane deformations sufficient for budding even for small *ε*. Conversely, for low ligand numbers and low binding strengths, there exist no particle designs that can bud in, although their performance can improve with evolutionary time.

In the regime of intermediate ligand numbers and binding strengths the ligand arrangement is crucial for the nanoparticle’s budding performance. This is the regime we focus on here. We define this “intermediate region” such that between 10% and 80% of the population penetrated the membrane. This corresponds to the mean total summed affinity of ϵ = 248.5*kT* and a standard deviation of 28.8*kT*, as shown in boxed region in Fig. 2 (b). We selected the particles from this region and removed duplicates, which left us with a population of 60260 unique particle designs. Interestingly, within the narrow intermediate range of total adhesion energy, nanoparticles with less active ligands and correspondingly higher individual ligand-membrane affinities cross the membrane more readily than particles with high ligand numbers. These results point to the importance of ligands arrangement for efficient particle budding.

## Structural Characterisation

Several representative examples of fit and unfit design are given in Fig. 3 (a). The difference between good and poorly performing particles is visible by eye, with the fit particles showing linearly conencted ligands, while poorely performing particles have ligands disconnected from each other or well clustered.

**Fig. 3.**
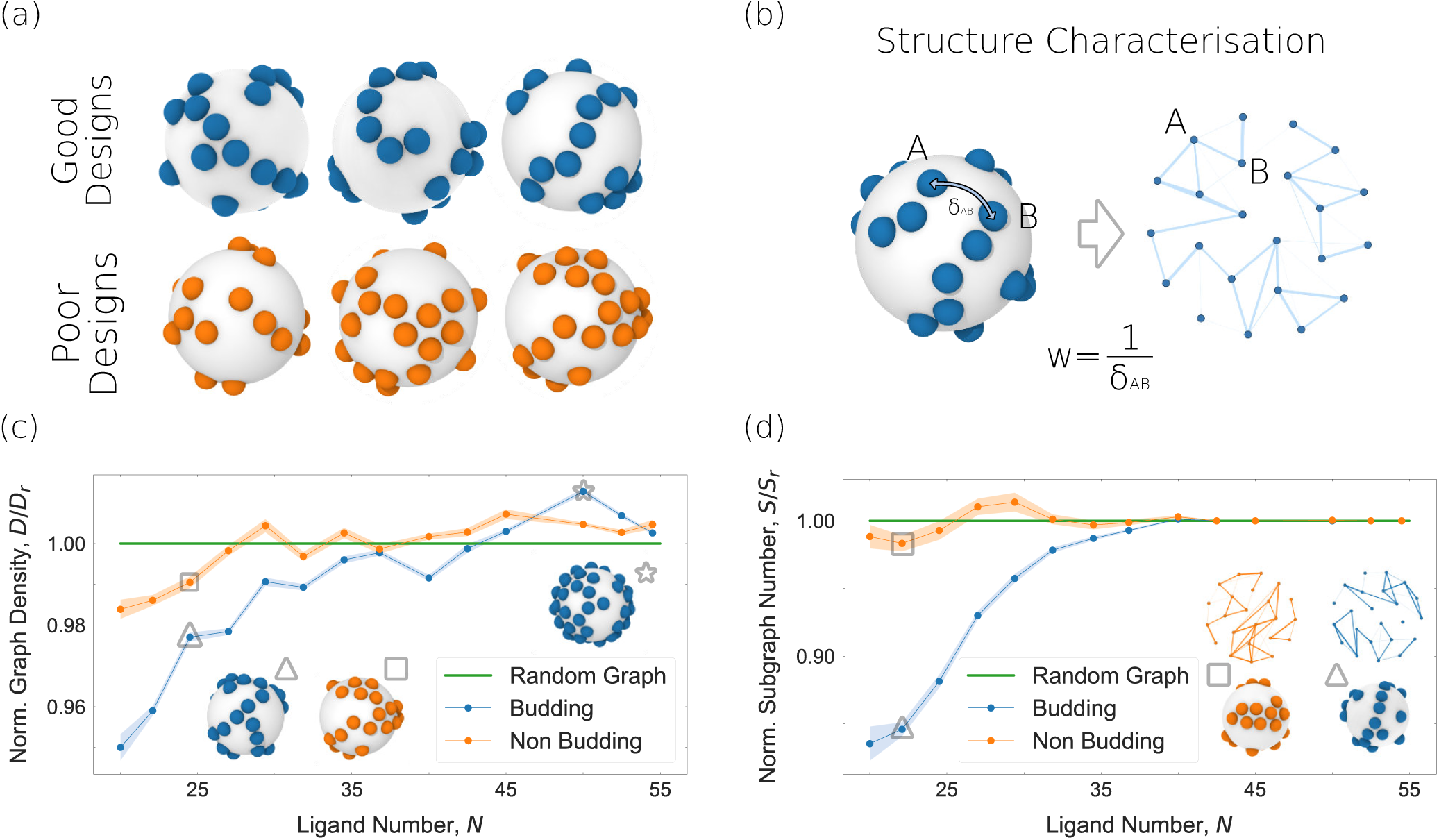
Structural analysis of the evolved particles. (a) Examples of evolved nanoparticle designs that exhibit good and poor performances. (b) The patterns of ligands on the particle are converted into a network, where the network edge weights are the reciprocal of the great arc distance between two ligands *δ*. This is repeated for all pairs of ligands with a distance *δ <* 3.3*σ*. Such a network is here visualised by a Fruchterman-Reingold projection(7). (c) The average normalised graph density for the budding and non-budding particles across the whole population. The insets show sample particles in each regime. (d) The average normalised subgraph size of budding and non-budding populations. The inset networks are two samples of budding and non-budding particles from the *N* = 23, *ϵ* = 10.0 *kT* dataset, each with 3 subgraphs; the differences in size distribution and the average connectivity of the subgraphs is visible to the eye.

To characterise nanoparticle designs in an unbiased way, we needed a quantitative measure to asses their structure. To this end, we translated the ligand positions on the surface of the particle into nodes of a network. The weights of the edges between the nodes are set to the inverse of the great arc distance between the ligands, as represented in Fig. 3 (b). This method of graph construction maps the surface of the particles in a rotationally invariant way. Since the closer ligands are to one another, the greater the weight of the edge connecting them, the network can be thought of as encoding the likelihood of cooperative binding between nodes. Once networks were constructed, they were pruned by removing edges which were above a threshold of 3.3*/σ*_0_, *σ*_0_ being the MD unit length, which effectively retains only nearest and next-to-nearest (second order) neighbours.

We found that two topological graph properties act as good structural descriptors that distinguish between successful and unsuccessful designs: the graph density *D* and the number of disconnected subgraphs *S*. The graph density of particle *i d*_*i*_ is defined as 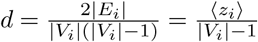, where |*E*_*i*_| and |*V*_*i*_| are the number of the particle network’s edges and vertices, respectively, and ⟨*z*_*i*_ ⟩ is the average degree of all nodes in the network. The graph density represents the ratio of the number of edges to the number of possible edges in an equivalent complete graph, and can be understood as a measure of how clustered and connected the points in the network are: a low network density corresponds to one with large ligand spacing, while a high density represents greater ligand clustering. The number of disconnected subgraphs in a particle *i, s*_*i*_, is number of separate regions of a graph that share no common nodes and indicates the spatial connectivity of ligands across the whole nanoparticle; in general, the lower the number of subgraphs the higher the connectivity for a fixed number of vertices. The population averages of these properties were taken across constant values of *N* to produce the mean density *D* and subgraph number *S* of designs at constant *N*. The same can be calculated for random particle designs at constant *N* (which have not undergone evolution), giving the mean density *D*_*r*_ and mean subgraph number *S*_*r*_.

## Identifying successful designs

The average normalised graph density *D/D*_*r*_ and subgraph number *S/S*_*r*_ for both the successful and unsuccessful population are shown in Fig. 3 (c) and (d), where *D*_*r*_ and *S*_*r*_ are the corresponding quantities for a uniformly random graph. The insets in Fig. 3 (c) and (d) illustrate a representative particles from each respective population.

The main structural difference between the successful and non-successful designs are best visible for low and intermediate ligand numbers, in our case below *N* = 35, where the number of possible variations in design is the largest. As shown in Fig. 3 (c), successful designs tend to have lower graph densities, meaning that the ligands have a lower average number of neighbours at given number of active ligands *N*. In contrast, Fig. 3 (d) clearly indicates that such particles also have lower average subgraph numbers. Therefore ligands in successful particles tend to be connected with one another by only few edges, covering a large angular range across the nanoparticle surface. Taken together, this indicates that successful designs are characterised by patterns in which each ligand is connected to all other ligands via chains, like those visible in Fig.3 (a) and in the inset in Fig.3 (c). Conversely, isolated “patches” of ligands perform very poorly (Fig.3 (a)). Interestingly, designs that have ligands uniformly distributed across the nanoparticle also show poor performance, with budding times on average 30% longer than that of the evolved succesful designs.

## Explaining successful designs

In searching for a physical explanation as to why the long chains of ligands exhibit superior membrane-crossing properties we noticed that unsuccessful designs, in which the ligands are clustered into disconnected patches, end up deforming the membrane but never becoming fully wrapped by it.

To better quantify this behaviour we measured the average rotational mean squared displacement Δ*θ*^2^ = ⟨ [*θ* (*τ*) – *θ* (0)]^2^⟩ as particles meet the membrane, bind to it, and deform it across the population of budding and non-budding particles. *θ* (*τ*) is the angle at time *τ* between a predefined nanoparticle axis and a predefined vector in the simulation box. The time evolution of the rotational displacement is presented in Fig. 4 (a). While at the beginning of the simulation the rotational displacement does not differ much between budding and non-budding particles, a significant discrepancy appears as the simulation progresses. Prior to budding, successful particles exhibit much larger rotational freedom compared to the non-budding particles. This rotational freedom enables particles to explore the transitional states needed to be able to wrap themselves in the membrane. Indeed, the frequency of budding events shown in Fig. 4 (a) illustrates the relative increase in the rotational freedom of budding particles with respect to non budding particles correlates and how this correlates with the proportion of the population that have budded at various times. As a control, we checked that if the rotational displacement is measured only until the point at which a particle buds, as opposed to for its full run length *t*_*run*_, the difference between the budding and non-budding population remains.

**Fig. 4.**
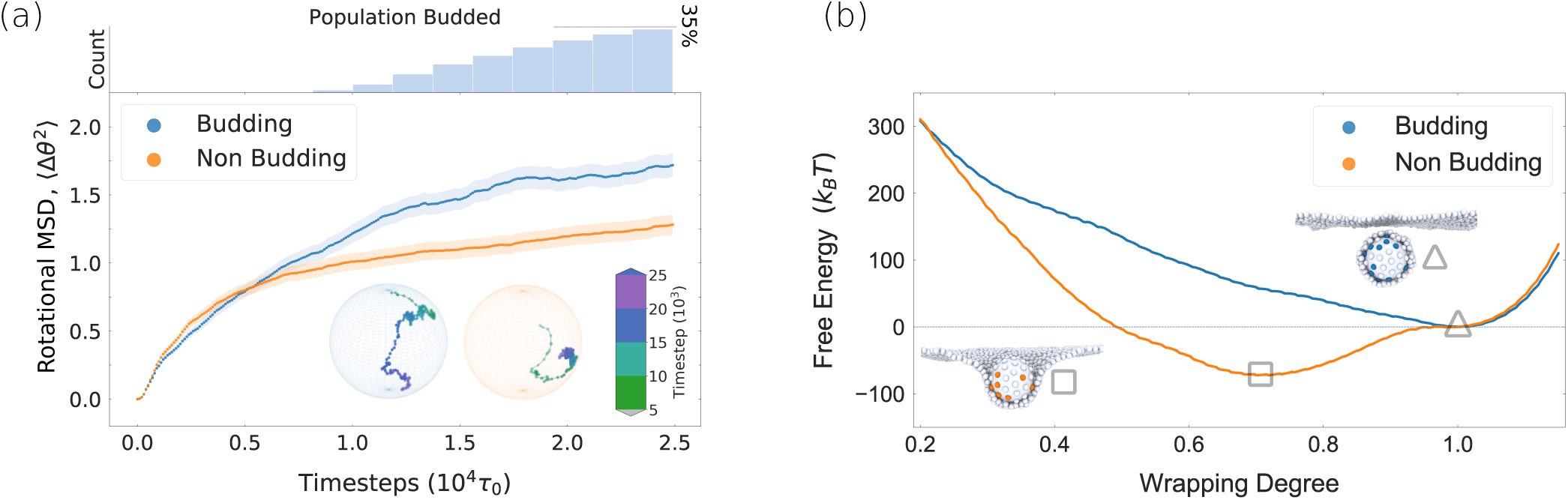
Explaining successful designs. (a) The rotational mean square displacement of the budding and non-budding population, shown for *ϵ* = 11*kT*, *N* = 22. The error bars represent the standard error of the mean for the ensemble averaged populations. The inset shows a sample rotational trajectory for one ligand on one particle from each respective subpopulation, while the top panel shows the count of the budding events. (b) Free energy as a function of a membrane wrapping for sample particles from the budding and non-budding populations (*ϵ* = 11*kT*, *N* = 22 in both cases). The insets show each particle at their representative membrane wrappings: the non-budding particle reaches the free energy minimum at the *≈* 75% membrane wrapping that is separated from full wrapping by a large barrier, while the budding particle reaches the free energy minimum at full wrapping.

This difference in the rotational mobility of budding and nonbudding particles implies that the purpose of the chains of ligands is to minimise the free energy barrier required for the particle wrapping. Fig. 4 (b) shows the free energy profile for budding for an example of a fit and unfit design, which carry the same total adhesion energy, computed using umbrella sampling with the degree of membrane wrapping as the reaction coordinate. Interestingly, the free energy minimum for the unfit particle is positioned at ∼75% membrane wrapping and is separated from the full wrapping by a large barrier. The free energy profile for the fit particle on the other hand reaches the global free energy minimum at full wrapping, without encountering any significant energy barriers en route. As this makes clear, evolution selects for designs that exhibit the lowest free energy barrier for membrane wrapping.

## Discussion and conclusions

By combining molecular dynamics simulations with genetic algorithms, we have been able to evolve nanostructures optimised for a specific function, selected for the ability to bud across the cell membrane. Even in this seemingly simple case, this approach revealed novel design rules that would not have been easy to guess, even for a well trained modeller.

At intermediate and low ligand numbers, the MD-GA scheme identified the nanoparticle designs that have a low free energy barrier for membrane crossing. Such designs are characterised by long chains of ligands, where the chains rarely cross or branch, but which effectively cover the particle leaving few gaps. One can think of these ligand chains acting like zippers, making membrane wrapping more reliable. Chains were often found together with small patches of ligands. While it is possible that such occasional patches have a functional role, this may instead be due to the limited number of evolutionary cycles.

The superior performance of the structures is based purely on the kinetics, since all the explored (*N, ε*) designs carry the same total adhesion energy. Indeed, for any practical application, the rate of membrane crossing is the key factor, rather than its thermodynamics. We therefore believe that the design rules identified here can aid the design of artificial nanoparticles for cell delivery, as well as possibly explain ligands patterns found in nature. For instance, low-density lipoprotein particles appear to carry membrane-binding protein arranged into long lines that envelop the particles (8). Analogously, membrane-binding proteins on some viruses display linear chain arrangements (9–11).

Previous studies incorporated evolutionary strategies within computer simulations mainly for the purpose of energy minimisation (12–14) or optimisation of interaction parameters (15, 16). Only few studies used such a combined approach to evolve a structure for a specific function. At a larger scale, Kriegman et al (17) used evolutionary algorithm within a physical engine environment to optimise for organisim design of a desired locomotion. At the nanoscale, Srinivasan et al. (18) combined an evolutionary algorithm with free energy calculations to reverse-design sequences on DNA-grafted colloids for a target colloidal crystal structure. This approach was based on identifying the free energy minimum, neglecting the kinetic effects that will be of crucial importance in an experimental realisation. The closest in spirit to the one presented here is the study by Miskin et al. (19), who combined covariance matrix adaptation evolution strategy with molecular dynamics simulations to evolve a particle shape that produces granular packing of a desired mechanical response.

All the previous MD-GA studies at the nanoscale focused on systems at high packing fractions, where computer simulations are relatively expensive, which somewhat limits the application of the combined scheme. Here we show that this combined approach might pose a far higher promise in the regime of low volume fractions, where functional nanostructures naturally operate.

While the previous body of work on the subject of nanoparticle uptake identified the importance of the nanoparticle shape (20–22) and ligand distribution(23), each of the studies explored only a small phase space of possible designs (∼10). Here we effectively explored tens of thousands of possible unique designs, and have identified novel, superior, design rules that previous studies did not consider. Indeed, the ability to efficiently explore a large phase-space of possible designs, in an unbiased way, is the main advantage of the computational approach presented here. We envision that the approach developed here can be of great help in identifying the design principles of a range nanostructures and nanomachines, such as protein filaments, lattices, and other higher-order structures, both in the context of biological and engineered systems.

We acknowledge support from EPSRC (JCF), MRC (BB and AS), the ERC grant NEPA (JK and AS), the Royal Society (AS), the UK Materials and Molecular Modelling Hub for computational resources, which is partially funded by EPSRC (EP/P020194/1).

## Methods

### Simulation Model

Simulations were carried out using the open source molecular dynamics package LAMMPS (4). The interactions between the active particle ligands and membrane beads were modelled using a cut-and-shifted Lennard-Jones potential, which takes the following form for the interparticle distances below the cut-off *r*_*c*_ = 1.8*σ*_0_ (*σ*_0_ being the MD unit of lengths) and is zero otherwise:

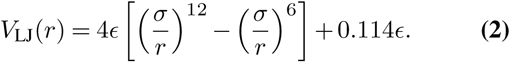

Here ϵ is the depth of the potential, representing the effective ligand-membrane interaction strength, and *σ* = *σ*_0_ is the contact distance between a ligand and a membrane bead.

The discretised positions of the ligands on the particle surface are set by an approximate sphere packing solution(24). The membrane was modelled using the single particle thick model developed by Yuan et al. (3) which is capable of fission and fusion events, with membrane parameters chosen to encode for a flat membrane with a bending rigidity of 15*kT*. Following the notation from the original paper (3), we chose the parameters 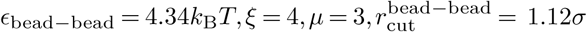, where *σ* = *σ*_0_ is also the membrane bead diameter. Simulations were run using a Langevin thermostat (with temperature set to 1.0 and damping parameter set to 1.0) within the *npH* ensemble for *t*_*run*_ = 25000 time steps, each of a size 0.01*τ*_0_, *τ*_0_ being the MD unit of time. The membrane contained 2900 particles with initial in plane dimensions 50*σ*_0_ by 50*σ*_0_, and a 200*σ*_0_ high box. The membrane centre of mass was tethered to the centre of the box by a spring. Particle coverage and budding was detected using a spatial clustering algorithm in LAMMPS, with the time point at which a significantly sized (*>* 30) group of membrane particles break away from the membrane bulk (along with the nanoparticle) being recorded as the time of budding.

### Particle Evolution

The genetic algorithm was built in-house using the DEAP python library(25) and used as a wrapper for LAMMPS simulations. The fitness function used in the genetic algorithm was chosen to drive greater particle coverage and drive faster budding (Eq. (1)). The rewards were set to reflect this, with the reward for budding *R*_*b*_ = 100, for coverage *R*_*c*_ = 1, and for budding time *R*_*t*_ = 10. Unique particle designs were tested 4 times under different initial rotational positions.

Each *ϵ, N* pair had its own evolutionary algorithm instance which was split into 4 subpopulations (demes) each with 20 individuals which ran independently in parallel during each generation until all individuals within them were assigned fitness scores and crossover mutation and cloning had occurred. At this point a single individual was chosen at random to migrate to another deme and replace the lowest fitness particle in that population (with the probability of being selected determined by its fitness normalised by the sum of all fitnesses in the deme) and the next generation proceeded in the same fashion. Selection tournaments were performed on 3 randomly selected particles, with the top particle in the tournament being allowed to survive and tournaments repeated until the population has been exhausted. Each instance of the algorithm was run for a total of 35 generations.

Parameters for the evolutionary algorithm were chosen to discourage premature fixation (the entire population becoming identical). The probability of crossover *P*_cross_ was set to 0.5, probability of mutation *P*_mut_ was set to 0.2 the independent probability of a genome bit being selected for swapping *P*_mutind_ was set to 0.15. Any gaps left after crossover and mutation are filled with clones of selected individuals. A tabulated form of these parameters is included in Table S1.

### Analysis

For the structural analysis of the ligand arrangements, graph edge weights were encoded using the reciprocal of the great arc distance between each pair of ligands on the surface. Weight cutoff for graph pruning was chosen to include (on average) first and second order nearest neighbours. Rotational mean square displacement (Rotational MSD) was calculated by taking the original position of an arbitrary ligand at timestep 0 and tracking the position of that same ligand over the course of the simulation run.

### Free energy calculation

To obtain the free energy profile as a function of the degree of nanoparticle wrapping umbrella simulations were carried out using the colvars module of LAMMPS.

The degree of wrapping *γ*_wrap_ is defined through the coordination number between the central carrier particle and the 2900 membrane particles. Using the reaction coordinate *γ*_wrap_ the simulation trajectories are biased by the harmonic potential 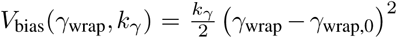, where *γ*_wrap,0_ denotes the position of the umbrella window and *k*_*γ*_ the force constant of the biasing potential.

During umbrella simulations 60 consecutive biasing windows are set by choosing *γ*_wrap,0_ in the interval [50, … 450] with *k*_*γ*_ = 0.5 *kT*. In each window the centre-of-mass position of the nanoparticle is sampled for 5000 time steps at a frequency 0.01 and each full simulation is repeated 200 times to obtain statistics. The resulting histograms for the individual windows are combined subsequently using the weighted histogram analysis method.

## Supplementary Note 1: Genetic algorithm

**Fig. 5.**
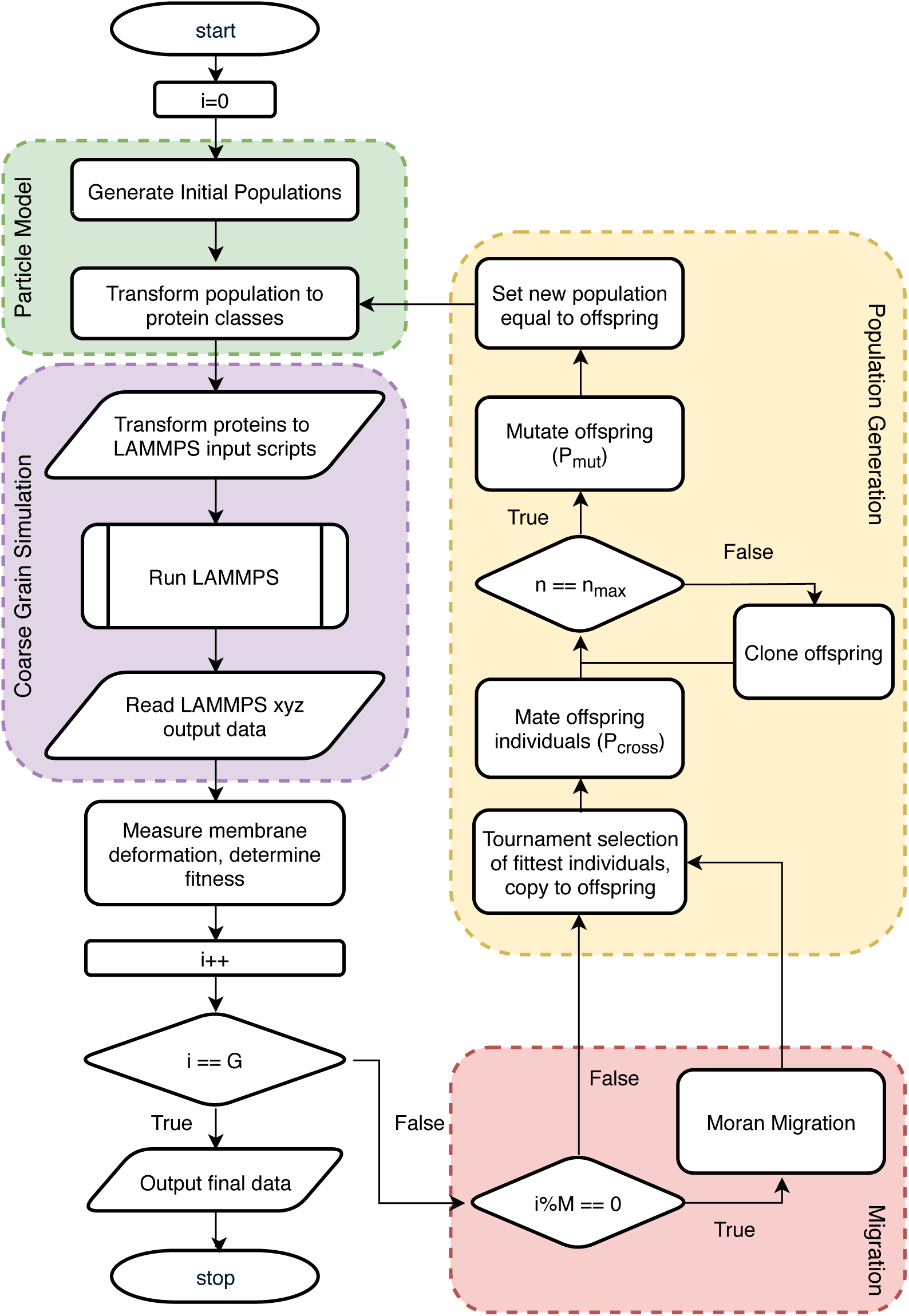
A program flowchart of the key steps of the genetic algorithm implemented in the study. Here *i* represents the current generation number and is iterated every time the loop is run. *G* is the number of generations that the GA is allowed to run for, in this study *G* = 35. *M* is the frequency of migration, in this study *M* = 1, so a migration took place every generation as described in the text. *n* is the population size and *n*_max_ is the target population size, in this case *n*_max_ = 20 for each deme. *P*_cross_ and *P*_mut_ are the probabilities of crossover and mutation occuring, as described in the main text. Here an *A* == *B* represents a test for equivalence between *A* and *B*, and “*A*%*B*” is the modulus operator which finds the whole number remainder of *A* divided by *B*.

**Table 1.** Parameters of the evolutionary algorithm and the simulations run as part of it. *R*_*b*_, *R*_*t*_ and *R*_*c*_ are the rewards for budding, speed of entry and coverage respectively. These are used in the calculation of fitness seen in equation 1 in the main text. *P*_cross_ and *P*_mut_ are the independent probabilities that an individual will be selected for crossover or mutation. If an individual is selected for crossover a different individual is selected at random with uniform probability from the rest of the deme population. *P*_mutind_ is the independent probability that each bit will be chosen to be swapped with another in the genome of an individual chosen for mutation, for each bit chosen a new position is chosen at random from the rest of the bits in the genome with uniform probability. *N*_gen_ is the number of generations, *n*_demes_ is the number of separated subpopulations, with *n*_max_ being the number of individuals in each deme. The total population of the entire system is *n*_demes_ *× n*_max_, for a total of 80 in this case. *M* is the number of individuals chosen for migration at the end of each generation. *n*_tour_ is the number of individuals selected for each round of tournament selection. *t*_run_ is the total runtime of each simulation, and *R* is the number of randomly oriented repeat simulations run for each new individual.

| *Parameter* | *Value* |
| --- | --- |
| $R_b$ | 100 |
| $R_t$ | 10 |
| $R_c$ | 1 |
| $P_{\text{cross}}$ | 0.5 |
| $P_{\text{mut}}$ | 0.2 |
| $P_{\text{mutind}}$ | 0.15 |
| $G$ | 35 |
| $n_{\text{demes}}$ | 4 |
| $n_{\text{max}}$ | 20 |
| $M$ | 1 |
| $n_{\text{tourn}}$ | 3 |
| $t_{\text{run}}$ | 25000 |
| $R$ | 4 |

